# Adult influence on juvenile phenotypes by stage-specific pheromone production

**DOI:** 10.1101/291591

**Authors:** Michael S. Werner, Marc H. Claaßen, Tess Renahan, Mohannad Dardiry, Ralf. J. Sommer

**Author notes:** These authors contributed equally (Michael S. Werner, Marc H. Claaßen, Tess Renahan).

## Abstract

Many animal and plant species respond to high or low population densities by phenotypic plasticity. To investigate if specific age classes and/or cross-generational signaling affect(s) phenotypic plasticity, we developed a dye-based method to differentiate co-occurring nematode age classes. We applied this method to *Pristionchus pacificus*, which develops a predatory mouth form to exploit alternative resources and kill competitors in response to high population densities. Remarkably, only adult, but not juvenile, crowding induces the predatory morph in other juveniles. Profiling of secreted metabolites throughout development with HPLC-MS combined with genetic mutants traced this result to the production of adult-specific pheromones. Specifically, the *P. pacificus*-specific di-ascaroside#1 that induces the predatory morph exhibits a binary induction in adults, even though mouth form is no longer plastic in adults. This cross-generational signaling between adults and juveniles may serve as an indication of rapidly increasing population size. Thus, phenotypic plasticity depends on critical age classes.

## Introduction

Population density is an important ecological parameter, with higher densities corresponding to increased competition for resources (Hastings, 2013). In addition to density-dependent selection (MacArthur, 1962; Travis et al., 2013), which operates on evolutionary time scales, some organisms can respond dynamically to population density through phenotypic plasticity. For example, plants can sense crowding by detecting the ratio of red (chlorophyll absorbing) to far red (non-absorbing) light, and respond by producing higher shoots (Dudley and Schmitt, 2015). Locusts undergo solitary to swarm (i.e. gregarious) transition, and aphids can develop wings, both as results of increased physical contact (Pener and Simpson, 2009; Simpson et al., 2001; Slogget and Weisser, 2004). Intriguingly, population density can also have cross-generational effects. For example, adult crowding of the desert locust *Schistocerca gregaria* (Maeno and Tanaka, 2008; Simpson and Miller, 2007) and migratory locust *Locusta migratoria* (Chen et al., 2015; Hamouda et al.) also influences the egg size, number, and morphology of their progeny; and high population densities of red squirrels elicit hormonal regulation in mothers to influence faster-developing offspring (Ben Dantzer et al., 2013). In many species, population density and cross-generational signaling are detected through pheromones, however the precise nature, mechanisms of induction, age-specificity, and exact ecological role are not well understood.

Nematodes are a powerful model system to investigate the mechanisms of density-dependent plasticity because many small molecule pheromones that affect plastic phenotypes have been characterized (Butcher, 2017; Butcher et al., 2007; Reuss et al., 2012). For example, in the model organism *Caenorhabditis elegans*, high population densities induce entry into a stress-resistant dormant ‘dauer’ stage (Fielenbach and Antebi, 2008). The decision to enter dauer was revealed to be regulated by a family of small molecule nematode-derived modular metabolites (NDMMs) called ascarosides that act as pheromones (Butcher et al., 2007; 2008; Jeong et al., 2005). Ascarosides consist of an ascarylose sugar with a fatty acid side chain and modular head and terminus groups (Figure 1A). The level and composition of ascarosides were later shown to be dependent on sex (Chasnov et al., 2007; Izrayelit et al., 2012) and development (Kaplan et al., 2011), although it is thought that dauer can be induced by all developmental stages (Golden and Riddle, 1982). Subsequent studies revealed that specific NDMMs also regulate other life history traits, such as mating (Chasnov et al., 2007; Izrayelit et al., 2012), social behavior (Srinivasan et al., 2012) and developmental speed (Ludewig et al., 2017). Although NDMMs are broadly conserved (Choe et al., 2012; Dong et al., 2018; Markov et al., 2016), inter- and intraspecific competition have driven the evolution of distinct response regimes for the same phenotypes (Bose et al., 2014; Choe et al., 2012; Diaz et al., 2014; Falcke et al., 2018; Greene et al., 2016). In addition, more complex structures have been observed that affect distinct plastic phenotypes (Bose et al., 2012).

**Figure 1.**
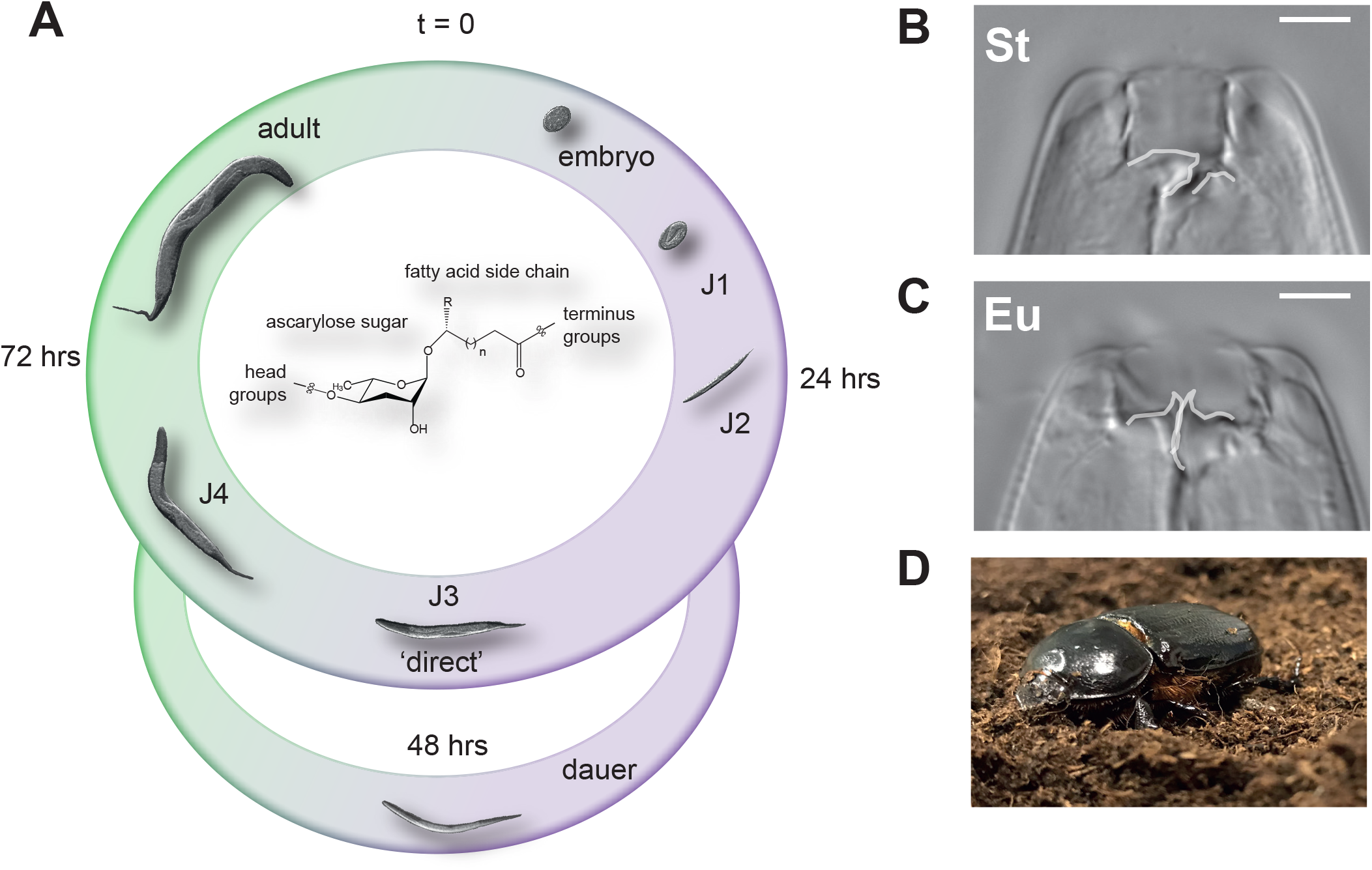
Life cycle and developmental plasticity of the model nematode *Pristionchus pacificus*. (A) The life cycle of *P. pacificus* consists of four juvenile stages (J1-4) until sexual maturation (adults). Like many nematodes *P. pacificus* can enter a long-living ‘dormant’ dauer state that is resistant to harsh environmental conditions. The decision to continue through the direct life cycle or enter dauer is regulated by small molecule excreted ascarosides (chemical structure adapted from (Butcher, 2017)). (B) *P. pacificus* can also adopt one of two possible feeding structures; either a microbivorous narrow-mouth (stenostomatous, St), or (C) an omnivorous wide-mouth (eurystomatous, Eu) with an extra tooth that can be utilized to kill and eat other nematodes or fungi. White lines indicate the presence of an extra tooth (right side) in the Eu morph or its absence in the St morph, and the dorsal tooth (left side), which is flint-like in St and hook-like in Eu. White scale bar indicates 5 µM. (D), *P. pacificus* is often found in a necromenic association with beetles (ex. shown here *Oryctes borbonicus*, photo taken by Tess Renahan) in the dauer state, and resumes the free living life cycle upon beetle death to feed on the ensuing microbial bloom.

In *Pristionchus pacificus*, a soil-associated nematode that is reliably found on scarab beetles (Figure 1A)(Herrmann et al., 2006; 2007; Sommer and McGaughran, 2013), an ascaroside dimer (dasc#1) that is not found in *C. elegans* regulates the development of a predatory mouth form (Bento et al., 2010a; Bose et al., 2012; Sommer et al., 2017). Mouth-form plasticity represents an example of a morphological novelty that results in predatory behavior to exploit additional resources and kill competitors. Specifically, adult *P. pacificus* exhibit either a narrow stenostomatous (St) mouth (Figure 1B), which is restricted to bacterial feeding, or a wide eurystomatous (Eu) mouth with an extra denticle (Figure 1C), which allows for feeding on bacteria and fungi (Sanghvi et al., 2016), and predation on other nematodes (Wilecki et al., 2015). This type of phenotypic plasticity is distinct from direct *vs*. indirect (dauer) development because it results in two alternative life history strategies in the adult (for review see Sommer & Mayer, 2015). Recent studies in *P. pacificus* have begun to investigate the dynamics and succession of nematodes on decomposing beetle carcasses to better understand the ecological significance of mouth-form plasticity (Meyer et al., 2017). These studies revealed that on a carcass (Figure 1D), *P. pacificus* exits the dauer diapause to feed on microbes, and then reenters dauer after food sources have been exhausted, displaying a ‘boom-and-bust’ ecology (Meyer et al., 2017; Sommer and McGaughran, 2013). Presumably different stages of this succession comprise different ratios of juveniles and adults, and recognizing the age-structure of a population as a juvenile could provide predictive value for adulthood. However, it is unknown whether the mouth-form decision is sensitive to crowding by different age classes. More broadly, while age classes are known to be important for population growth and density-dependent selection {Hastings:2013dn, Charlesworth:1994ww, Charlesworth:1970ks}, their role in phenotypic plasticity has thus far been largely unexplored.

While nematodes have many experimental advantages, including easy laboratory culture and advanced genetic, genomic, and the aforementioned chemical tools, their small size has made investigations at the organismal level and in experimental ecology challenging. For example, no *in vivo* methodologies are currently available to label distinct populations without the need for transgenics, which is only available in select model organisms such as *C. elegans*, *P. pacificus*, and some of their relatives. Here, we combine a novel dye-staining method with the first developmental pheromone profiling in *P. pacificus* to study potential effects of age on density-dependent plasticity. This vital-dye method allows tracking adults with juveniles, or juveniles with juveniles, and can be applied to any nematode system that can be cultured under laboratory conditions. In contrast to dauer, we found that mouth form is strongly affected by cross-generational signalling. Specifically, only adult crowding induces the predatory morph, which is controlled by adult-specific pheromones.

## Results

### A vital dye method for labeling nematode populations

To directly test if different age groups of nematodes influence plastic phenotypes, we required two synchronized populations to co-habit the same space, yet still be able to identify worms from different age groups. To do so, we developed a dye-staining methodology to robustly differentiate between nematode populations. After trying several vital dyes, we identified that Neutral Red (Thomas and Lana, 2008) and CellTracker Green BODIPY (Thermo) stain nematode intestines brightly and specifically to their respective channels (Figures 2A-E and S1). These dyes stain all nematodes tested including *C. elegans* (Figure S2) and dauer larvae (Figure S3A,B). They also last more than three days (Figure S3C-G), allowing long-term tracking of mixed nematode populations. Importantly, neither Neutral Red nor CellTracker Green staining affects viability, developmental rate, or the formation of specific morphological structures, such as *P. pacificus* mouth form (Figure S4). Thus, Neutral Red and CellTracker Green allow specific labeling of worm populations to study age-dependent effects on phenotypes.

**Figure 2.**
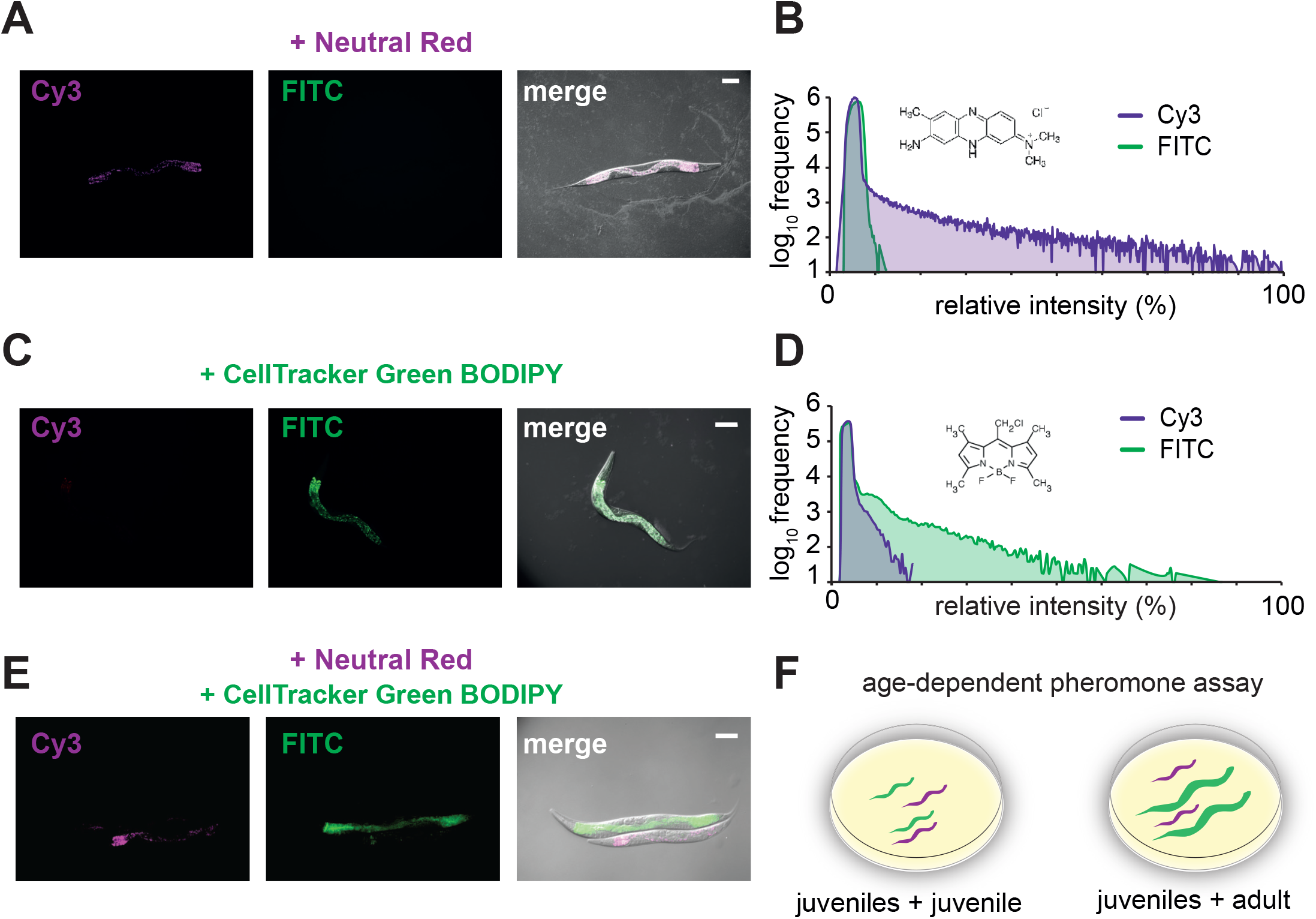
Vital-dye method in nematodes allows mixing different populations together. (AE) *P. pacificus* were stained with either 0.005% Neutral Red or 50 µM CellTracker Green Bodipy (Thermo) and viewed using Cy3 and FITC filters. Images were merged with Differential interference contrast (DIC), scale bar = 100 µM. An example of relative intensities of each fluorescence channel are displayed in the histograms (right) with the chemical structure of Neutral Red or CellTracker Green Bodipy. (F) Age-dependent functional pheromone assay: Experimental juveniles were stained with Neutral Red, and challenged with CellTracker Green Bodipy-stained juveniles or adults on standard condition NGM agar plates seeded with 300 µl OP50 *E. coli*. Three days later, only red-positive and green-negative adults were phenotyped.

### Adult but not juvenile crowding induces the predatory mouth form in *P. pacificus*

To assess potential intra- or inter-generational influence on *P. pacificus* mouth form we stained juveniles of the highly St strain RSC017 with Neutral Red, and added an increasing number of CellTracker Green-stained RSC017 adults or juveniles (Figure 2F, 3A). Three days later we phenotyped red animals that had developed into adults, but showed no green staining. To ascertain potential differences between adding juveniles or adults, we performed a binomial regression on Eu count data from multiple independent biological replicates (n>3), with age and number of individuals added as fixed effects (Transparent Methods, Table S1). We observed a significant increase in Eu worms in response to adults, but not juveniles (*p*=2.59 x10^-2^; for display summed percents are shown in Figure 3B,C). Almost half (48%) of the population developed the Eu mouth form with just 500 adult animals, which is a greater than 50-fold induction compared to side-by-side controls (Figure 3B,C). We were also curious if dauers, which have a thickened cuticle and represent a distinct stage in the boom-and-bust life cycle of nematodes, could still respond to adults. Indeed, the same trend that was observed with juveniles was seen with dauers (*p*=2.96×10^−3^), albeit to a more muted extent (Figure 3D,E). Specifically, with a total of 200 dauers and 500 adults, 25.7% of dauers become Eu, whereas only 1.8% of dauers become Eu on a plate containing 700 dauers (and no adults) (Figure 3D). Collectively, these data indicate that adult crowding specifically induces the Eu mouth form.

**Figure 3.**
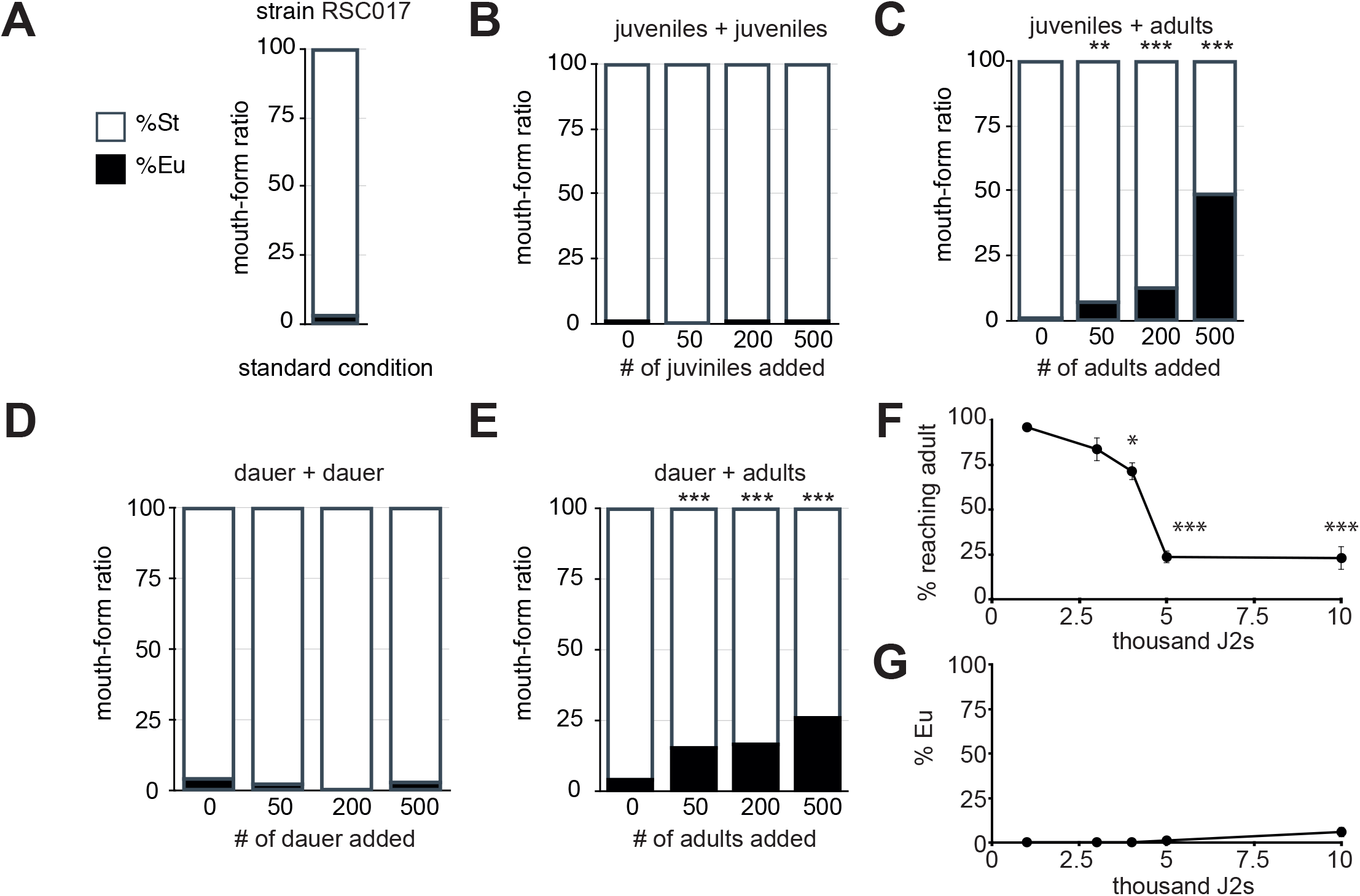
Vital-dye method confirms adult-specific density effect on mouth form. (A) The wild isolate RSC017 grown in standard conditions (5 young adults passed to fresh plates, progeny phenotyped 4 days later) are highly stenostomatous (<10%, *n*=102). (B,C) Mouth form ratios of Neutral Red-stained J2s, and (D,E) dauers, with increasing number of CellTracker Green-stained competitors, as described in Figure 2 (n=3-5 independent biological replicates per experiment, with total n>100 per experiment). Overall significance between strain and age was determined by a binomial linear regression (see Transparent Methods), and pairwise comparisons were assessed by Fisher’s exact test on summed Eu counts (Significance codes: ‘***’ < 0.001, ‘**’ <0.01, ‘*’ <0.05). Mouth forms were phenotyped at 40-100x on a Zeiss Axio Imager 2 light microscope. (F) Percent reaching adulthood and percent Eu of those that reached adulthood (G) after increasing numbers of J2s are added to standard 6 cm NGM agar plates with 300 µl OP50 *E.coli* bacteria (n=2 biological replicates, with total *n*>200 for percent reaching adulthood, and total n>100 for mouth form. Significance was determined by a binomial regression).

Even though we did not detect a mouth-form switch in large populations of J2s or dauers, and food was still visible on plates containing the most animals (500 adults and 200 juveniles), we could not completely rule out the possible effect of food availability on mouth form. As a proxy for starvation, we conducted assays with greatly increased numbers of juveniles from 1,000 to 10,000 that would rapidly deplete bacterial food. We noticed a stark cliff in the fraction of juveniles that reach adulthood at 4,000-5,000 animals, arguing that food is a limiting resource at this population density (Figure 3F). Importantly however, in these plates we still did not see a shift in mouth form (Figure 3G) (p=0.99, binomial regression, Table S1). With an overwhelming 10,000 worms on a plate, 5.8% were Eu, compared to 48% in the presence of only 500 adults. While longer-term starvation may yet have an impact on mouth form, under our experimental conditions it appears to be negligible.

### Adult but not juvenile secretions induce the Eu mouth form

As mouth-form plasticity in *P. pacificus* is regulated by nematode-derived modular metabolites (NDMMs)(Bose et al., 2012), we wondered if the difference between adults and juveniles resulted from differences in secreted NDMMs. To test this hypothesis we added secretions from adult or juvenile worms to RSC017 (highly St) juveniles. We found that adult secretions from both the laboratory stain RS2333 (highly Eu) and RSC017 led to a significant increase in the Eu morph relative to juvenile secretions (*p*=5.27×10^−06^, 1.33×10^−3^, respectively, Fisher’s exact test)(Figure 4). To confirm the effect was caused by ascaroside pheromones, we exposed RSC017 juveniles to supernatant from a *daf-22.1;22.2* double mutant, which exhibits virtually no ascaroside production in both *C. elegans* and *P. pacificus* (Golden and Riddle, 1985; Markov et al., 2016). Again, juvenile secretion had no impact on Eu frequency, but in contrast to wild-type supernatants, we observed no significant increase in Eu frequency with adult secretions (*p*=0.8324, Fisher’s exact test, Figure 4). Thus, adult-specific NDMMs induce development of the Eu mouth form.

**Figure 4.**
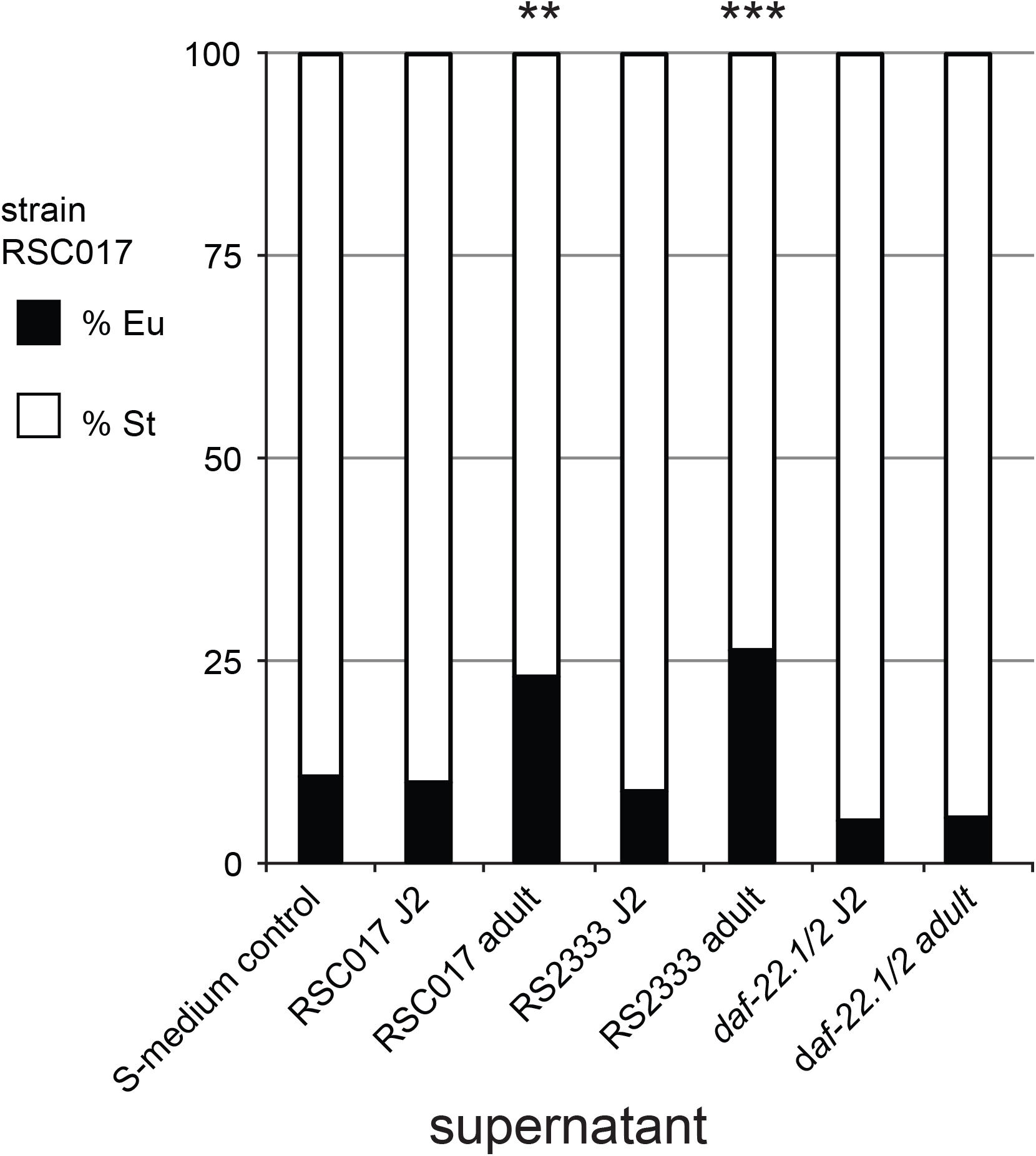
Adult-specific secretions induce predatory morph in juveniles. Highly St strain RSC017 juveniles were exposed to J2 and adult supernatants of its own strain, and to the J2 and adult supernatants of highly Eu strain RS2333. Mouth form was phenotyped three days later. Worms exposed to J2 secretions remained highly St, while worms exposed to adult secretions had a small but significant increase in Eu morphs (p<0.05, Fisher’s exact test). Supernatants from the double mutant *daf-22.1/2*, which has deficient ascaroside pheromone production, did not elicit juvenile or adult increase in Eu. Worms exposed to the S-media control also remained highly St. n=4 independent biological replicates for RS2333 and *daf-22.1/2* secretions, and n=2 independent biological replicates for RSC017 adult and juvenile secretions, with an average count of 55 animals per replicate. For display, total Eu and St counts are presented as percentages (Significance codes: ‘***’ < 0.001, ‘**’ <0.01, ‘*’ <0.05).

### Developmental-staged NDMM profiles reveal adult-specific synthesis of dasc#1

Next, we investigated whether the difference between adult and juvenile pheromones is one of dosage, or of identity. To answer this question and verify age-specific differences in pheromones, we profiled *P. pacificus* NDMM levels in two strains and at three time points throughout development. We used RS2333 and RSC017 and measured the exo-metabolomes of juvenile stage 2 (J2s, 24 hrs), J3s (48 hrs) and J4/adults (72 hrs) from a constant culture with excess OP50 bacterial food (Figures 5A,B, S5, Materials and methods). To assess potential differences in pheromone levels we performed a linear regression with the area under the curve for each NDMM (aoc) (Figure S5) as the response variable. Stage and strain were modeled as fixed effects, and because we performed separate regression analyses for each pheromone, we adjusted the resulting *p* values for multiple testing using false discovery rate (FDR)(see Table S2 for *p* and *FDR* values between stage and strain). We observed that there was a significant affect of developmental stage on the levels of ascr#9, ascr#12, npar#1, and dasc#1, and that ubas#1 and #2 are strain and stage specific (FDR<0.05). Interestingly, dasc#1 is the most potent known Eu-inducing compound when tested as a single synthesized compound, while npar#1 is both Eu- and dauer-inducing (Figure 5C,D,F-I) (Bose et al., 2012). Closer inspection revealed dasc#1, npar#1, and ascr#9 increase throughout development in both strains, and dasc#1 peaks in adults in RS2333 (p<0.05, student’s two-tailed *t*-test between 72 hrs and 24 hrs for each NDMM in both strains, and also 72 hrs and 48 hrs for dasc#1 in RS233, Table S3). Intriguingly, the trajectory of dasc#1 appeared binary in both strains (Figure 5F,G). In fact our statistical model for dasc#1 fits better if we assume cubic rather than linear growth (*ΔAIC*=3.958). In contrast, ascr#9, which was also statistically up-regulated but does not affect known plastic phenotypes (Bose et al., 2012), displays a more gradual increase in both strains (Figure 5E,J,K), and the model fits better with linear growth (AIC_linear_ – AIC_cubic_= −1.208). Meanwhile, the trajectory of npar#1 appears strain-specific (Figure 5H,I). Hence the mode of induction is NDMM-specific, and the kinetics of production may be related to their roles in phenotypic plasticity.

**Figure 5.**
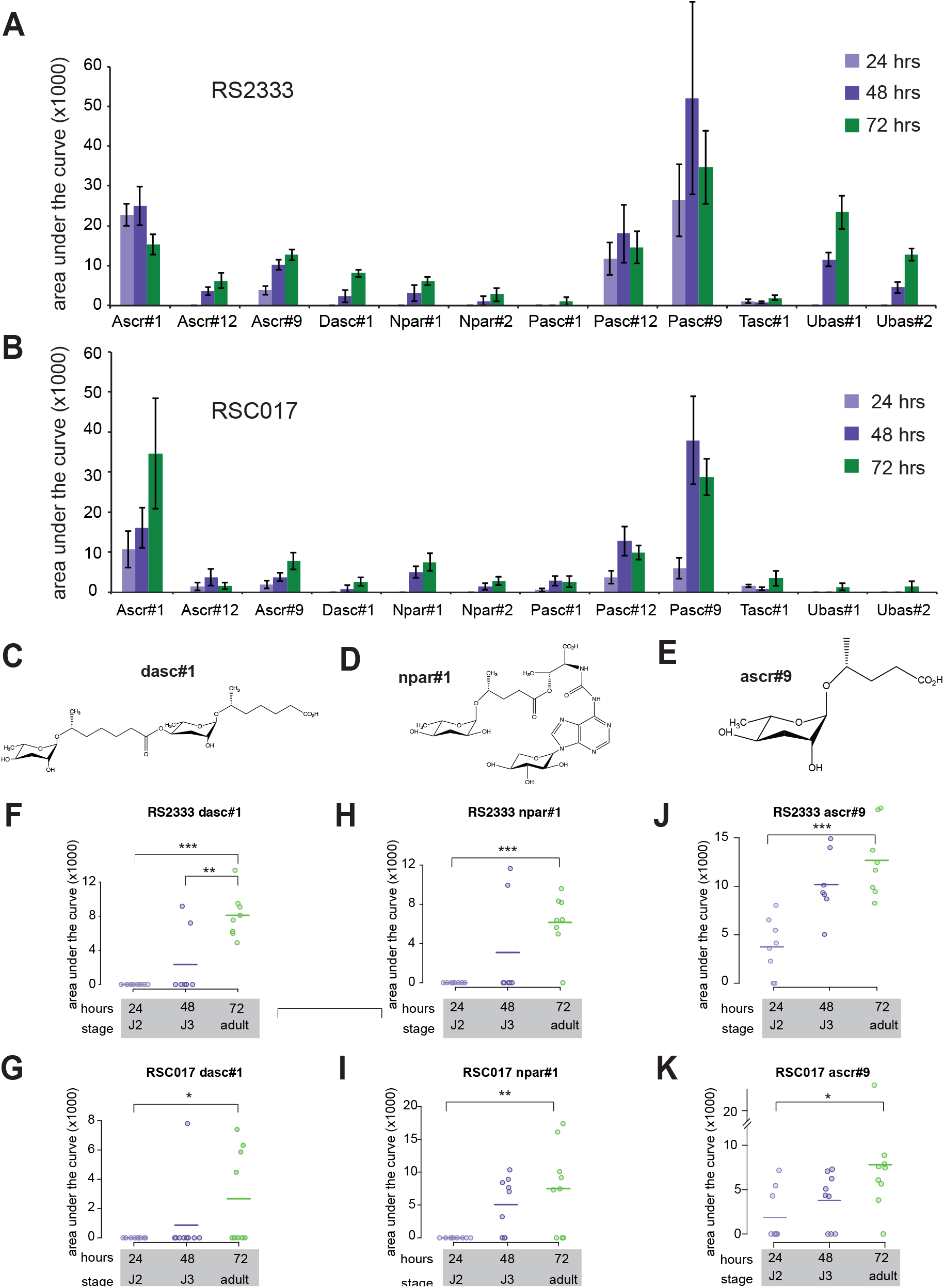
Time-resolved Nematode-Derived Modular Metabolites (NDMMs) in *Pristionchus pacificus*. (A) Time resolved secretion profile of nematode derived modular metabolites from the wild-type laboratory strain RS2333 and (B) wild isolate RSC017. Data is presented as the mean of 8 (RS2333) and 9 (RSC017) biological replicates, and error bars represent standard error of the mean (SEM). (C-E) Chemical structures of adult specific NDMMs dasc#1, npar#1, and ascr#9, as described in the Small Molecule Identifier Database (http://www.smid-db.org/), produced in ChemDraw. (F-K) Time-resolved abundance of dasc#1, npar#1, and ascr#9 NDMMs in RS2333 and RSC017. Each data point represents a biological replicate, and for comparison to (A-B) lines represent mean abundance. *P* values calculated by a 2-tailed students *t*-test (Significance codes: ‘***’ < 0.001, ‘**’ <0.01, ‘*’ <0.05).

In principle, the increase in abundance of certain pheromones could be a result of a concomitant increase in body mass, however several observations indicate more targeted regulation. First, no other compounds were significantly different in our linear model. Second, an analysis of previously published RNA-seq data (Baskaran et al., 2015) reveals the increase in NDMM abundance corresponds to an increase in transcription of the thiolase *Ppa-daf-22.1* (Figure S6), the most downstream enzyme in the β-oxidation pathway of ascaroside synthesis. Third, pasc#9 and pasc#12 actually exhibit a peak in abundance at the 48 hr/J3 time point, rather than in 72 hrs/adults. Finally, we profiled the endo-metabolome of eggs, and found appreciable amounts of ascr#1, #9, #12, and pasc#9, but little to no traces of other ascaroside derivatives (Figure S5C), suggesting age-specific synthesis rather than release. Together, these results suggest that the observed increase in ubas#1 and #2, ascr#9, npar#1, and dasc#1 over time corresponds to age-specific production. The observation that dasc#1 is produced specifically during the juvenile-to-adult transition is especially intriguing because adults are no longer able to switch mouth forms, hinting at cross-generational signaling.

## Discussion

Here, we introduce a novel dye-based method that allowed us to assess cross-generational influence on mouth form. Our results demonstrate adult crowding induces the Eu predatory morph, and that this effect is a result of age-specific pheromones. In doing so, we provide the first multi-stage time series of pheromone production in *P. pacificus*, which shows that dasc#1 exhibits a surprising ‘off-on’ switch-like induction pattern. Collectively, our results argue that adults represent a critical age group (Charlesworth, 1972) in nematode populations.

Our developmental profiling revealed an increase in two NDMMs that affect plastic phenotypes. The observation that this trend mirrors the transcriptional regulation of enzymes involved in NDMM synthesis argues that the stage-dependent increase is not simply a result of an increase in body mass, but rather that these molecules are programmed for stage-specific induction. The binary ‘off-on’ kinetics might reflect a population level feedback loop, such that the production of density-sensing pheromones is based on a threshold level of previously produced pheromones. It is also worth noting that while npar#1 is the major dauer-inducing pheromone in *P. pacificus* (Bose et al., 2012), we did not observe dauers in any experimental setup described herein. Thus, it seems that mouth-form phenotype is the first-level plastic response to population density. Presumably higher concentrations are required for dauer induction, reflecting a calculated response strategy depending on the level of crowding or duration of starvation. Interestingly, the effect of adult supernatants was noticeably less (23%-26% Eu) than of adult worms (up to 48% with only 500 adults). It is difficult to compare the amount of pheromone concentrations between experiments, but presumably worms in the vital-dye assay experienced a greater local concentration as they were in direct physical contact with each other, compared to worms in the supernatant assay.

Among the many environmental influences on mouth form (Werner et al., 2017), population density and starvation are perhaps the most ecologically relevant. However, teasing apart these two factors has proven difficult (Bento et al., 2010b). Here, we demonstrate that while a strong shift is observed with adult-specific pheromones, no such effect was seen under limited resource conditions. Thus, age-specific crowding is sufficient to induce the Eu mouth form. Nevertheless, this does not preclude that long-term starvation could also have an effect. Determining the relative contributions of these factors to mouth form will be important to better understand the sophisticated ecological response strategies of *P. pacificus*, nematodes, and phenotypic plasticity in general.

Why do adults and not juveniles affect mouth form? Given that St animals can develop faster (Serobyan et al., 2013), there may be a ‘race’ to sexual maturation in emergent populations at low densities. But as the nematode population increases, there will likely be a commensurate decrease in bacterial populations. When faced with competition from other nematodes, *P. pacificus* has a particular advantage in developing the Eu morph; their expanded dietary range includes their competition. Indeed, when nematode prey is the only available food source, the Eu morph provides longer life spans and more progeny than the St morph (Serobyan et al., 2014). When resources become depleted as population size increases, *C. elegans* and other monomorphic nematodes may enter dauer and disperse (Frézal and Félix, 2015). But in St-biased dimorphic strains of *P. pacificus*, juveniles may switch to the Eu morph in response to adults as a first-level indication of rapidly increasing population size (Figure 6). Then, after prolonged starvation and crowding, worms will presumably enter dauer. By analogy to economic models of population growth (Malthus et al., 1992; Trewavas, 2002) mouth-form plasticity is a ‘technological innovation’ to temporarily escape a Malthusian resource trap. To what extent this occurs in nature, or with *P. pacificus* strains that are highly Eu, remains to be determined.

**Figure 6.**
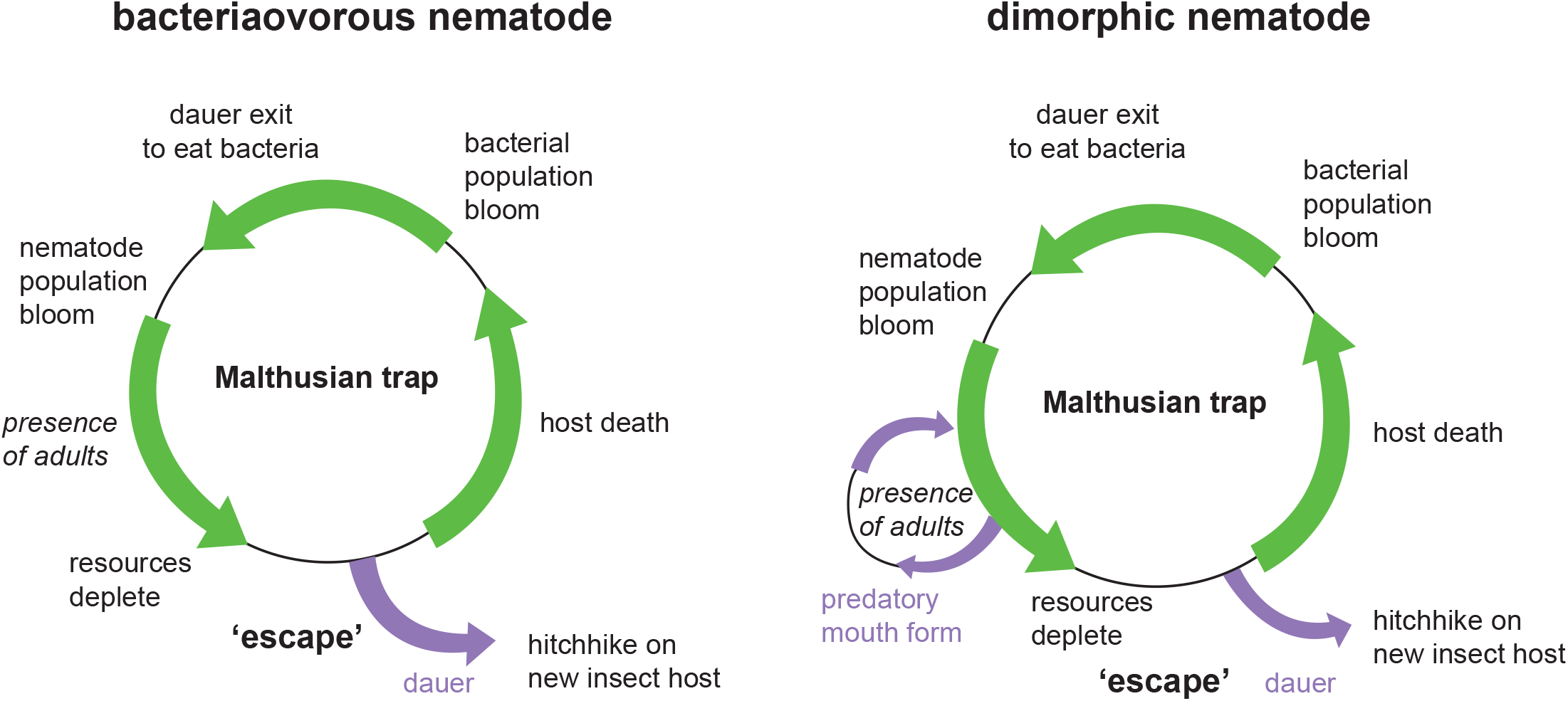
Conceptual model of the role of critical age classes in mouth-form phenotypic plasticity. Conceptual life cycle models of monomorphic or dimorphic mouth form nematodes. In an isolated niche such as a decaying insect carcass, at some point microbial food supplies will run out, leading to a Malthusian catastrophe. Nematodes escape this trap by entering the dauer state, hitchhiking to a new insect carrier, and re-starting the cycle. Dimorphic nematodes may sense the impending catastrophe earlier by recognizing an abundance of adults in the population, and switching to the Eu morph to exploit new resources and kill competitors. By analogy to economic models, the mouth form switch is a technological innovation to temporarily escape a Malthusian resource trap.

The evolution of dimorphic mouth forms is one among myriad nematode ecological strategies. For example, entomopathogenic nematodes release their symbiont bacteria in insect hosts to establish their preferred food source, and some release antibiotics to kill off competing bacteria and fungi from other entomopathogenic species (Griffin, 2012). Some free-living species, like those of the genus *Oscheius*, refrain from combat and stealthily feed and reproduce amidst warring entomopathogenic species. Interspecific killing also occurs in gonorchoristic species, in which both mated and virgin males are killed, implying fighting not just for mates but for resources as well (O’Callaghan et al., 2014; Zenner et al., 2014). Reproductive strategies also exist, and hermaphroditic species have an advantage over gonachristic species when colonizing a new niche, such as an insect carcass (Campos-Herrera, 2015). Meanwhile insect hosts and colonizing nematodes have their own distinct pheromone-based attraction and toxicity (Cinkornpumin et al.; Renahan and Hong, 2017). Finally, the renaissance of *C. elegans* sampling from around the world (Cook et al., 2017; Evans et al., 2016; Félix et al., 2013; Petersen et al., 2014; Poullet and Braendle, 2015) is rapidly building a resource of wild isolates that will almost certainly have different and fascinating ecologies. We hope our method for labeling and then combining different nematode populations on the same plate will aid in studies to identify these strategies. Perhaps the time is also ripe to complement these studies with more sophisticated ecological modelling that can lead to testable hypotheses.

Although beyond the scope of this manuscript, the cross-generational communication we observed could in principle reflect an intended signal from adults to juveniles, i.e. kin selection (Bourke, 2014). However, we favor a more simplistic view that juveniles have evolved to recognize adult-produced metabolites. Regardless of these interpretations, our results argue that age classes are a critical factor in density-dependent plasticity, as has been theorized in density-dependent selection (Charlesworth, 1994).

## Limitations of the Study

Given the ubiquity of certain traits in reproductive adults and their contribution to population growth, we suspect similar results will be found in other systems. However, it may depend on the phenotype and system being studied. For example, the population dynamics of nematodes (fast hermaphroditic reproduction) may be sufficiently different from other species such that our findings are not extendable in every case. In addition, our method of staining different populations, while fast and easy, is particular to nematodes.

## Methods

*See Transparent Methods in Supplemental Information

## Acknowledgements

We would like to thank all members of the Sommer lab, Dr. Talia Karasov, Dr. Hernan Burbano and Moises Exposito-Alonso for guidance with statistical analysis, and Dr. Adrian Striet (Max Planck Institute), and Dr. Cameron Weadick (University of Sussex) for thoughtful critique and discussion.

## Author Contributions

MSW and RJS conceived of the project. MC conducted pheromone profiling. MSW and TR designed and conducted dye-labeling experiments. TR and MC performed supernatant experiments. MD and MSW considered ecological implications. MSW and TR wrote the manuscript with input and edits from all authors.

## Supplemental Information

### Transparent Methods

#### Nematode strains and husbandry

*P. pacificus* Wild-type RS2333 (California) and RSC017 (La Réunion) strains were kept on 6 cm nematode growth media (NGM) plates seeded with OP50 and kept at 20°C. RSC017 is highly St and does not predate on other nematodes, and thus was used for biological assays instead of the highly Eu, predatory RS2333. To induce dauer, mixed-stage plates with little to no OP50 were washed with M9 and the resulting worm pellets were used in a modified ‘White Trap’ method. Worm pellets were placed on killed *Tenebrio molitor* grubs and dispersing dauers were collected in surrounding MilliQ water. Age of dauers ranged from one week to one month.

#### Dye staining

A stock solution of Neutral Red was prepared by dissolving 0.5 mg in 10 ml 5% acetic acid and stored at −20° C. Working solutions were prepared by 100x dilution in M9, aliquoted, stored at −20°C, and thawed directly before use. Working solutions were kept for approximately 1 month. Stock solutions of 10 mM Green Bodipy were made in DMSO and stored −20. J2s were prepared from 20-40 x 6 cm plates 6 days after passaging 5 worms to each plate on 300 µl OP50. Worms were washed from plates with M9 into a conical tube, and then filtered through 2×20 µM filters (Millipore) placed between rubber gaskets. The flow-through contains mostly J2 and some J3, which were pelleted by centrifugation, 8 seconds on a table-top eppdendorf centrifuge 5424, reaching approximately 10,000 x g. The juvenile pellet was then either re-suspended in 1 ml Neutral Red working solution, or in 1 ml M9 and split to two tubes, then recentrifuged, and then re-suspended in either 1 ml working solution Neutral Red (0.005% in M9) or 1 ml 50 µM Green BODIPY (Thermo) in M9. Tubes were then rotated for 3 hours in the dark, then washed by centrifugation as before, and re-suspended in 1 ml M9. This was repeated 3-4x until the dye was no longer visible in the worm pellet. Then, the concentration of worms was determined by aliquoting 2 µl onto a glass coverslip in 5 technical replicates, and counted under a dissecting microscope. Finally the appropriate number of animals was added to 6 cm plates that had been previously seeded with 300 µl OP50, and incubated at 20°C. After 3 days, 100% of worms exhibited Neutral Red staining (*n*=50, Supplementary figure 4). Dauers and J2s recovered after Neutral Red staining developed at the same developmental speed (3-4 days) and with the same mouth-form ratio as control worms recovered side-by-side (100% St for both, Supplementary figure 5, *n*=30). Dauers and J2s stained with Cell tracker Green BODIPY (50 µM) (Thermo) were similar, although less efficiently stained compared to Neutral Red. On day 4, 90% retained intestinal fluorescence (Supplementary figure 4), although brightness decreased with the number of days. Mouth-form ratios of dauers or J2s in +/- 50 µM Cell tracker Green BODIPY also developed at equivalent rates and mouth-form ratios (Supplementary figure 5). Lower than 25 µM did not yield strongly fluorescent worms after three hours. Cell Tracker Blue CMAC (Thermo) was also used at 50 µM and imaged 3 days post-staining for *P. pacificus*, and one day post-staining for *C. elegans*. However, due to the higher fluorescent background in the blue light spectrum in both *P. pacificus* and *C. elegans*, we performed all experiments using only Neutral Red and Cell tracker Green BODIPY.

#### Microscopy

All images were taken on a Zeiss Axio Imager 2 with an Axiocam 506 mono, and processed using Zen2 pro software. Image brightness and contrast were enhanced in ImageJ with a minimum displayed value of 10 and maximum of 100 for all images in Fig 2, and Supplementary figures 4 and 5, and a minimum of 21 and maximum of 117 for Supplementary figure 3. The following exposure times were used for all images: Cy3 (peak emission = 561, exposure = 80 ms), FITC (peak emission = 519, exposure = 150 ms), Dapi (peak emission = 465, exposure = 80 ms), DIC (exposure = 80-140 ms).

#### Mixed culture experiments and statistical analysis

We performed mixed culture experiment presented in figure 2 with 3 to 5 independent biological replicates, and a minimum total number of counts *n* > 100 (median counts per replicate for J2=29 and the median counts per replicate for dauers=27). J2 or dauers were stained with Neutral Red as described in the ‘Dye Staining’ method section, then added to green-stained J2, dauer, or adult populations on 6 cm plates with 300 µl OP50 and incubated at 20° C. To ensure consistent bacterial food supply, we added 1 ml more overnight OP50-LB to each plate on the following day, then air-dried under a chemical fume hood for 1 hour, then returned to 20° C. On days 3-4, we phenotyped ‘red’ adults that exhibited no ‘green’ staining. To assess whether the age of the ‘green’ surrounding population affects the mouth form of the dependent variable ‘red’ J2s we performed a binomial regression on Eu counts (i.e. “successes”) weighted by the number of counts per replicate and the number added as a fixed effect, using a generalized linear model from the standard statistical package in R:

glm(formula=cbind(Eu,total)~’stage_added’ * ‘#_added’, data=’J2/Da’, family="binomial"))

See Supplementary figure 6a for a table containing the resulting *p* values. The AIC for our models (78.97 for J2s and 89.59 for dauers) was substantially lower than the null hypothesis (220.16 for J2s and 147.29 for dauers), arguing a reasonable fit. For pair-wise comparisons of the effect of age for a given number of added animals, we performed a post-hoc Fisher’s exact test on a contingency table containing the summed counts of Eu and St observations. For display, we converted Eu counts into percent of total in figure 2, with the *p* values between the same number of animals added indicated over the adult-added population (Significance codes: 0 ‘***’ 0.001, ‘**’ 0.01, ‘*’ 0.05).

#### Measuring the effect of food depletion on mouth form

To verify that starvation was not a factor in our mixed culture experiments, we added increasing number of J2s to standard 6 cm plates with 300 µl OP50 to rapidly consume bacterial food, and measured both the amount of animals that reached adulthood, and the percent Eu in each population for two biological replicates. To assess the affects of added J2s to each dependent variable we performed a binomial regression with count data weighted by the total number of counts for each replicate:

glm(formula = cbind(reached_adult, total)~thousand_J2s, data=data_2, family="binomial"))

*p* values indicate a significant difference in percent reaching adult as a function of J2s added, but not in percent Eu (Table S1).

#### Supernatant collection and assays

Strains RS2333, RSC017, and RS2333-*daf-22.1;22.2* were raised in 10 mL liquid culture as in the time-resolved NDMM collections (see below). For each time point, 9 mL of the supernatant was lyophilized overnight, extracted again overnight with 90% ethanol (diluted in Millipore water) while being stirred, and centrifuged (4000g, 10 min, 4C). The solvent was evaporated and the solid re-dissolved with 1 mL Millipore water. This clear extract was then directly used for the assays. One mL of the supernatant was cleaned for HPLC-MS analysis (refer to pheromone profiling: HPLC-MS sample preparation) for quality control. For the assays, RSC017 was synchronized by bleaching and added to plates seeded with 300 µl OP50. The supernatants were added to the RSCO17 J2s in two 500 µl increments (for a total of 1ml supernatant) and dried for 30 minutes in a sterile hood after each addition. Plates were kept at 20C and adult mouth forms were screened three days later.

#### HPLC-MS sample preparation for normal exo-metabolome and time resolved analysis

To collect staged phermone profiles, we seeded 35 x 6 cm plates with 5 worms each, and bleached 5-6 days later when gravid to collect eggs/J1s. These were then seeded in 6 x 10 mL flasks with OP50 as described in Werner et al., 2017 (Werner et al., 2017). Then at 24, 48, or 72 hr time intervals, supernatant was obtained by centriguation (>4,000 x g, 4°C for 10 minutes). 1 mL supernatant was adsorbed onto a SPE-C8 cartridge (Thermo Scinetific Hypersept C8 100 mg/1mL), conditioned with 1 mL MeOH followed by 2 mL Millipore water. The adsorbed material was then washed with 200 uL water and subsequently eluted with 200 uL MeOH. This extract was then measured directly via HPLC-qTof MS (Bruker ImpactII).

#### HPLC-MS measurement

20 uL extract was injected into a Thermo ultimate 3000 HPLC equipped with a Sigma-Alderich Ascentis Express C18 2.7um 10mm x 4.6mm column at 20 °C with a flow of 500 uL/min. All MS measurements have been performed in negative ion mode and molecules are detected as [MH]^-^ Ions. The solvent gradient started with 5 % acetonitrile (ACN)/ 95 % water (both containing 0.1 % formic acid) for 2 min. After this equilibration step, the ACN proportion has been increased to 65 % over 8 min, then to 100 % ACN in 1.2 min followed by a hold step for 8.8 min. Afterwards, they system was flushed to 5 % ACN with 2 min equilibration for a total of 22 min. For calibration, a sodium formiat cluster building solution has been automatically injected in the first 2 minutes of each run. Data analysis was performed with TASQ version 1.0 from Bruker Daltonics. Extracted ion chromatograms for each well-known compound with a mass width of 0.1 m/z and time slices of 0.5 min around the expected retention time have been produced after calibrating and baseline correction. Assignment errors have been corrected with the provided MRSQ value.

#### Statistical analysis of NDMMs

NDMM levels were compared simultaneously between strains and developmental stages by a linear model in R: lm(‘NDMM’ ~ ‘developmental stage’ * ‘strain’, data=’data.frame’)). *P* values between stages and strains were adjusted for multiple testing by a false discovery rate correction. The level of fit between linear vs. exponential growth was determined by the Akaike information criterion (AIC). The lowest AIC for iterations of different exponents (n=1,2,3…) was used for comparison to the simple linear model. While significant in both cases, for consistency we present the original *p* values from the original linear model in Supplementary Figure 8.

### Supplemental Figure Legends

**Figure S1, related to Figure 2.**
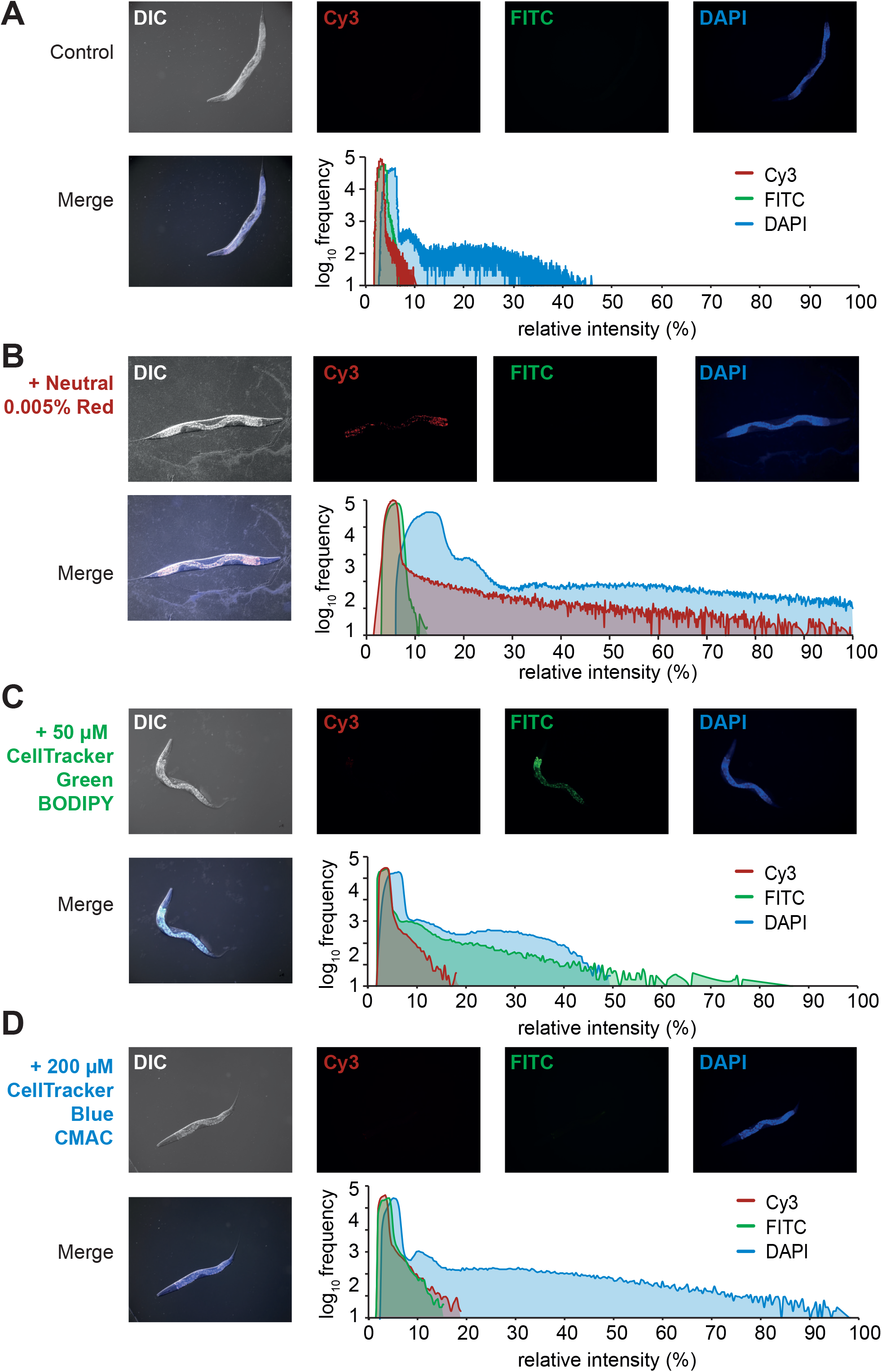
Vital dye staining of *Pristionchus pacificus*. (A) Control *P. pacificus* imaged with Cy3, FITC, and DAPI filters, and a merge with Differential Interference Contrast (DIC). Histogram on the right represents quantification of intensity with each filter. (B) Same as (A) but stained with 0.005% Neutral Red, (C), 50 µM CellTracker Green Bodipy (Thermo Fischer), or (D) 50 µM CellTracker Blue CMAC Dye (Thermo Fischer). J2s were stained (see Materials and Methods), and ensuing adult animals were imaged 3 days later on a Zeiss Axio Imager 2 with an Axiocam 506 mono, and processed using Zen2 pro software. Image brightness and contrast were enhanced in ImageJ for display, with a minimum displayed value of 10 and maximum of 100 for all images. Note that while Neutral Red and CellTracker Green staining are bright and specific to their respective channels, CellTracker Blue is indistinguishable from background fluorescence.

**Figure S2, related to Figure 2.**
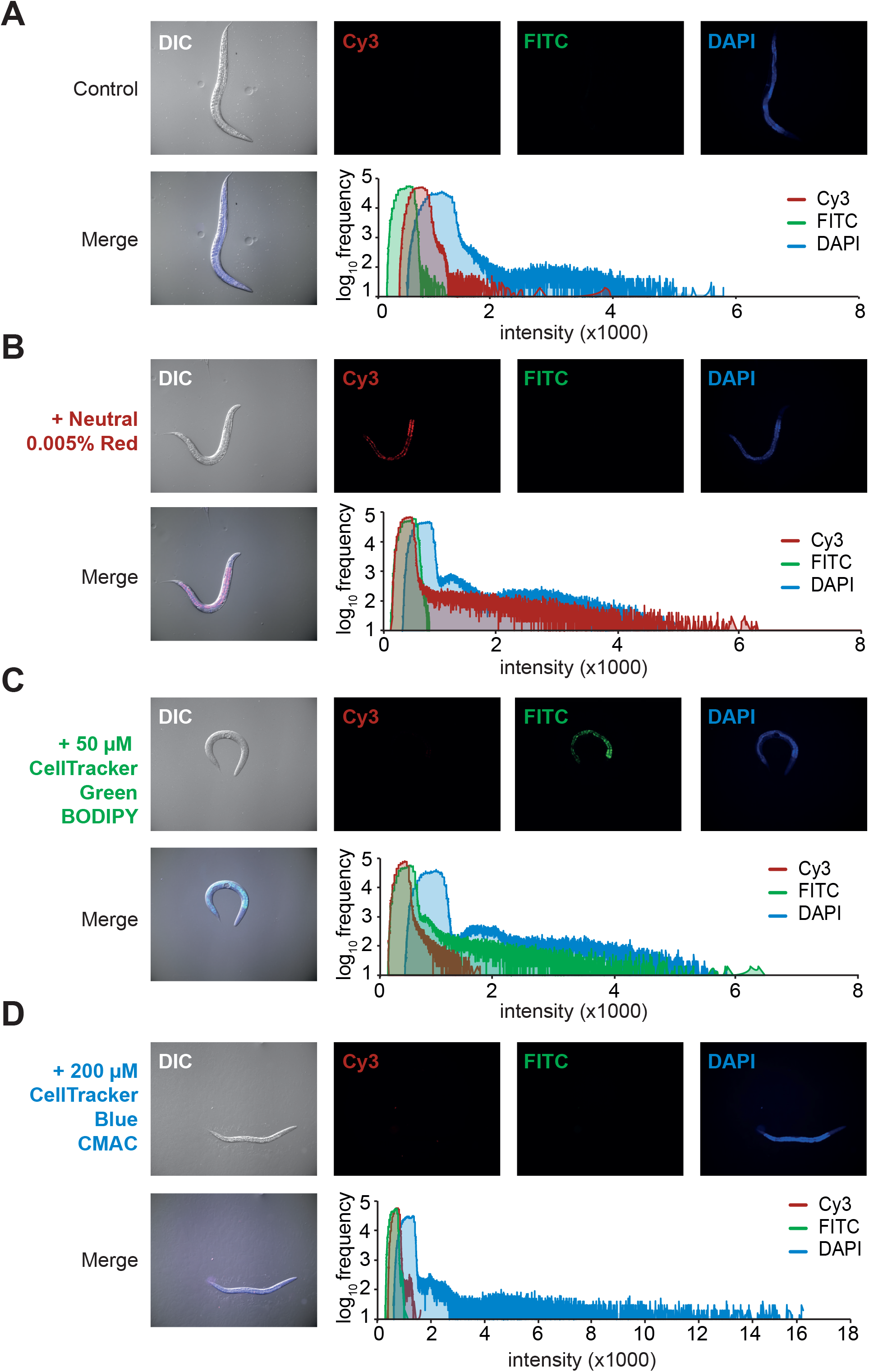
Vital dye staining of *Caenorhabditis elegans*. (A-D) Same as Supplementary Figure 1, but with *C. elegans*.

**Figure S3, related to Figure 2.**
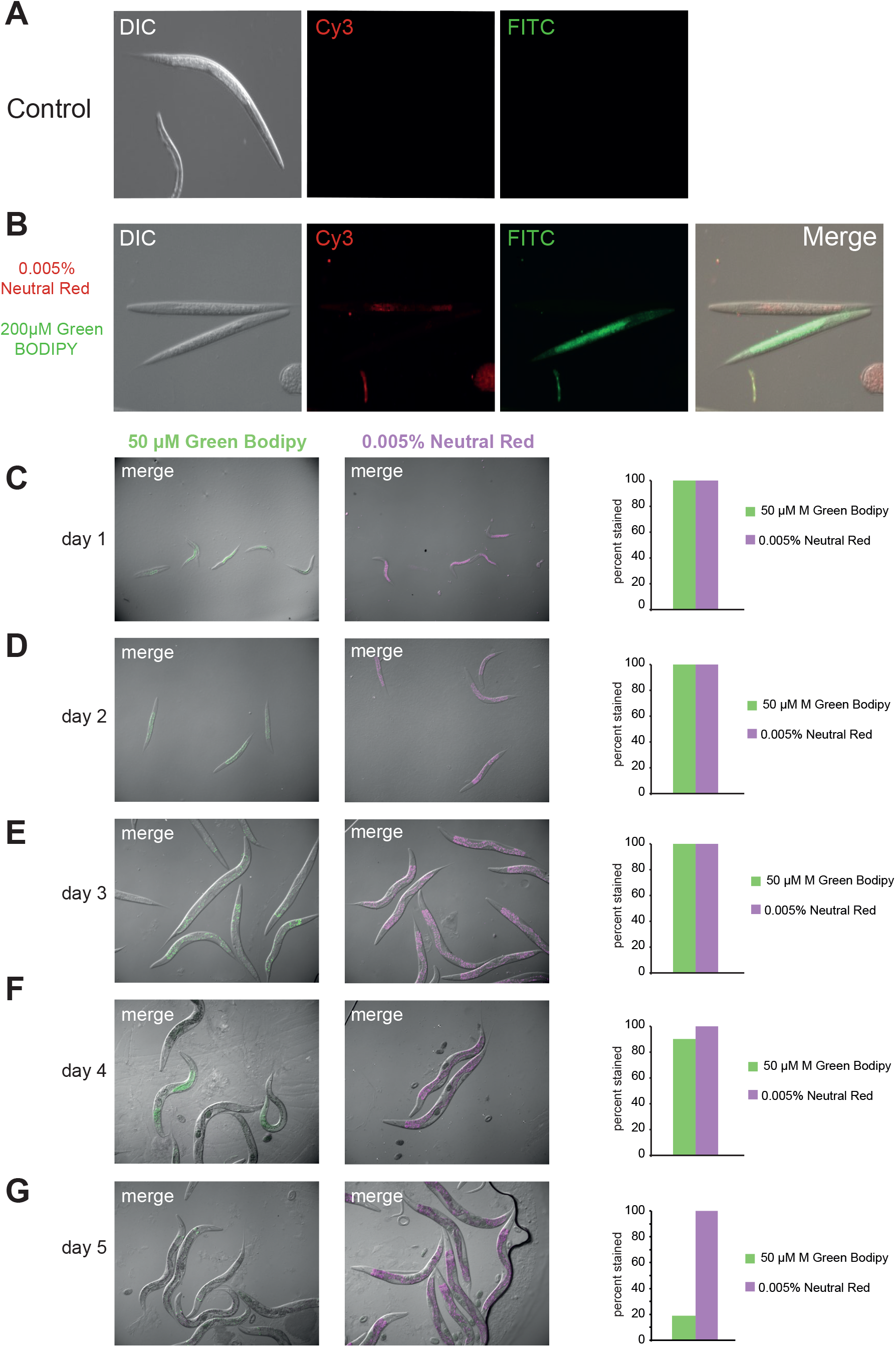
Vital dye staining of Pristionchus pacificus dauers, and duration of staining. (A) Control *P. pacificus* dauer imaged with DIC, Cy3, and FITC filters. (B) Dauers stained with either 0.005% Neutral Red or 50 µM CellTracker Green Bodipy and imaged immediately after staining with DIC, Cy3, and FITC filters and merged with DIC. Images were taken using Zeiss Axio Imager 2 with an Axiocam 506 mono, processed using Zen2pro software, and adjusted in ImageJ, with a display value minimum of 21 and maximum of 117. (C-G) 50 µM Cell Tracker Green Bodipy and 0.005% Neutral Red-stained J2s were imaged every day for five days. Percent of individuals retaining the dyes are shown in panels next to each microscope image for each day. Both stains are seen in all organisms for three days; Neutral Red (NR) persists for at least five, while the number of Green Bodipy (GB) –stained worms drops on day four. All images are merged with DIC, n=31 GB, 63 NR day 1, 68 GB, 56 NR day 2, 50 GB, 50 NR day 3, 50 GB, 50 NR day 4, 50 GB, 50 NR day 5.

**Figure S4, related to Figure 2.**
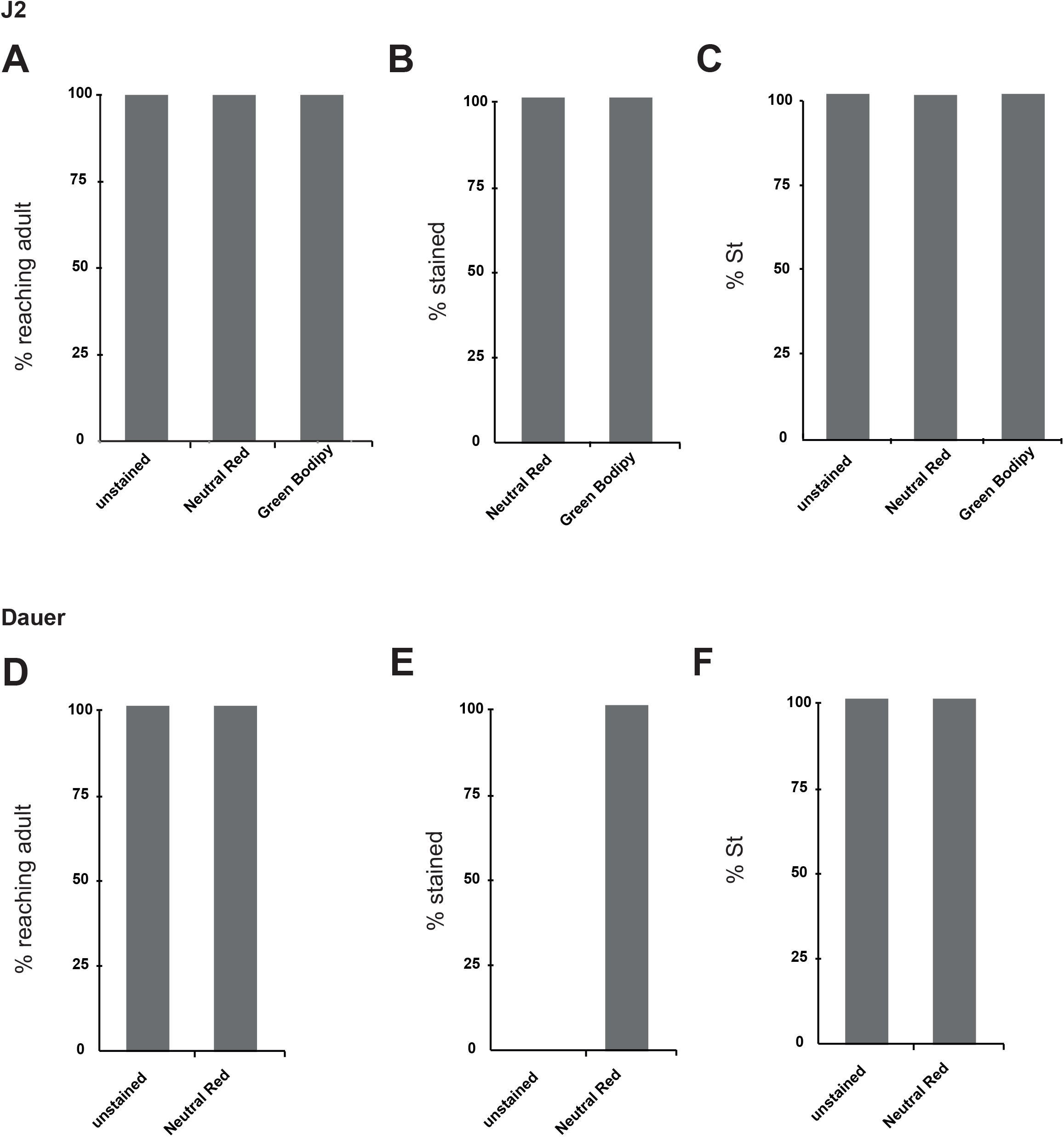
Vital dye staining does not affect *P. pacificus* mouth form or development. (A) Neutral Red and CellTracker Green Bodipy-stained J2s reach adulthood at the same rate as unstained J2s (3 days). (B) All of the J2s stained retain the dye in adulthood in the intestine. (C) Neither dye affects mouth form; both unstained and stained worms remain 100% St (n=30). (D-F) Same as for (A-C) except with dauers instead of J2s, and only with Neutral Red.

**Figure S5, related to Figure 5.**
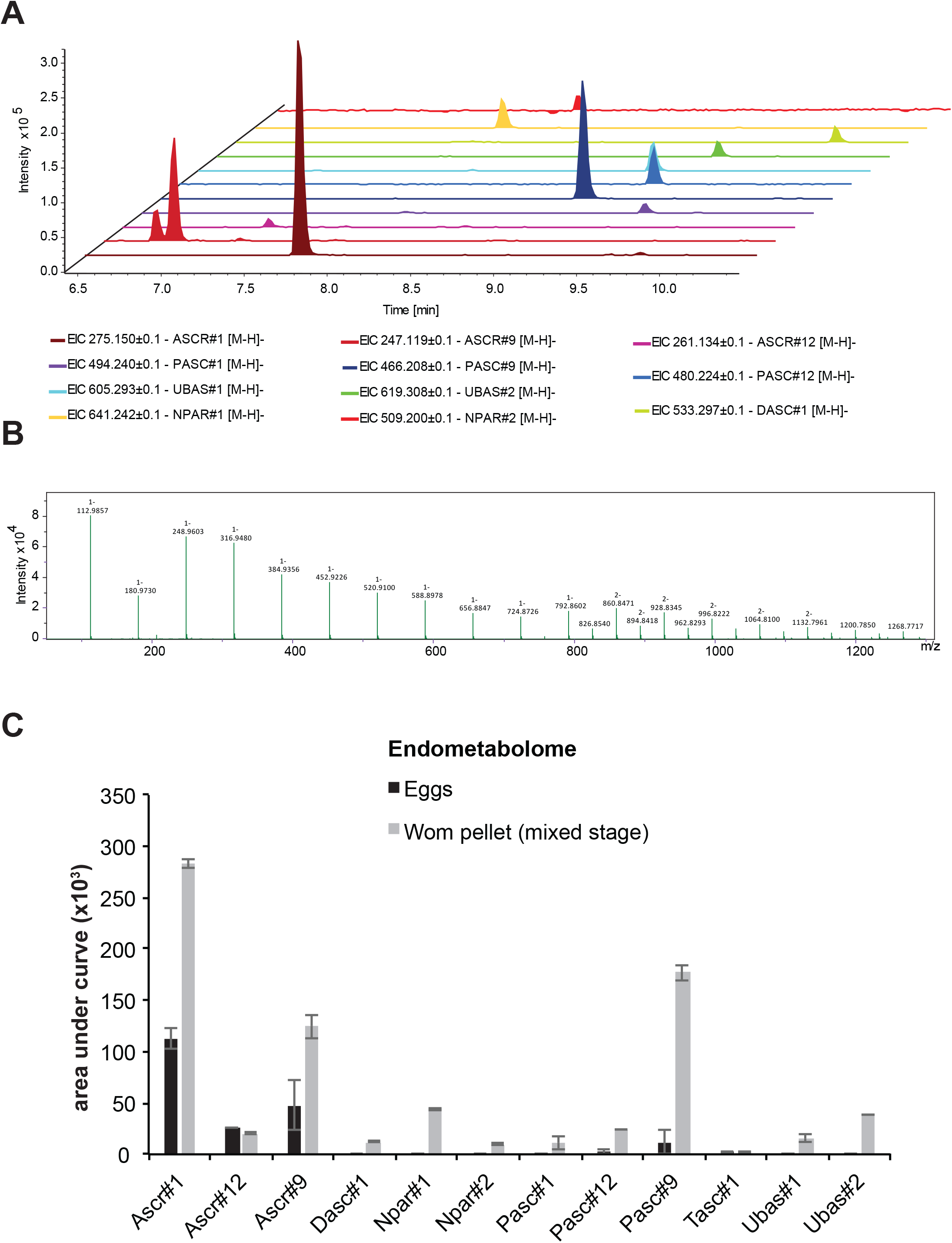
Pheromone-profiling quality control. (A) Extracted ion traces (width 0.1 m/z) of 11 of the 12 NDMMs used in this publication from a seven-day mixed-stage sample, double peak of 247.12 m/z indicate isomeric structures (Part#9/Ascr#9). (B) Example of an averaged spectrum over a calibration segment, sodium-formiat cluster building solution has been used to ensure high mass accuracy in each run. (C) Comparison of an endometabolome sample from a seven day mixed-stage cultured compared to the endometabolome of eggs, produced by using bleached eggs from 80 x 60 mm plates.

**Figure S6, related to Figure 5.**
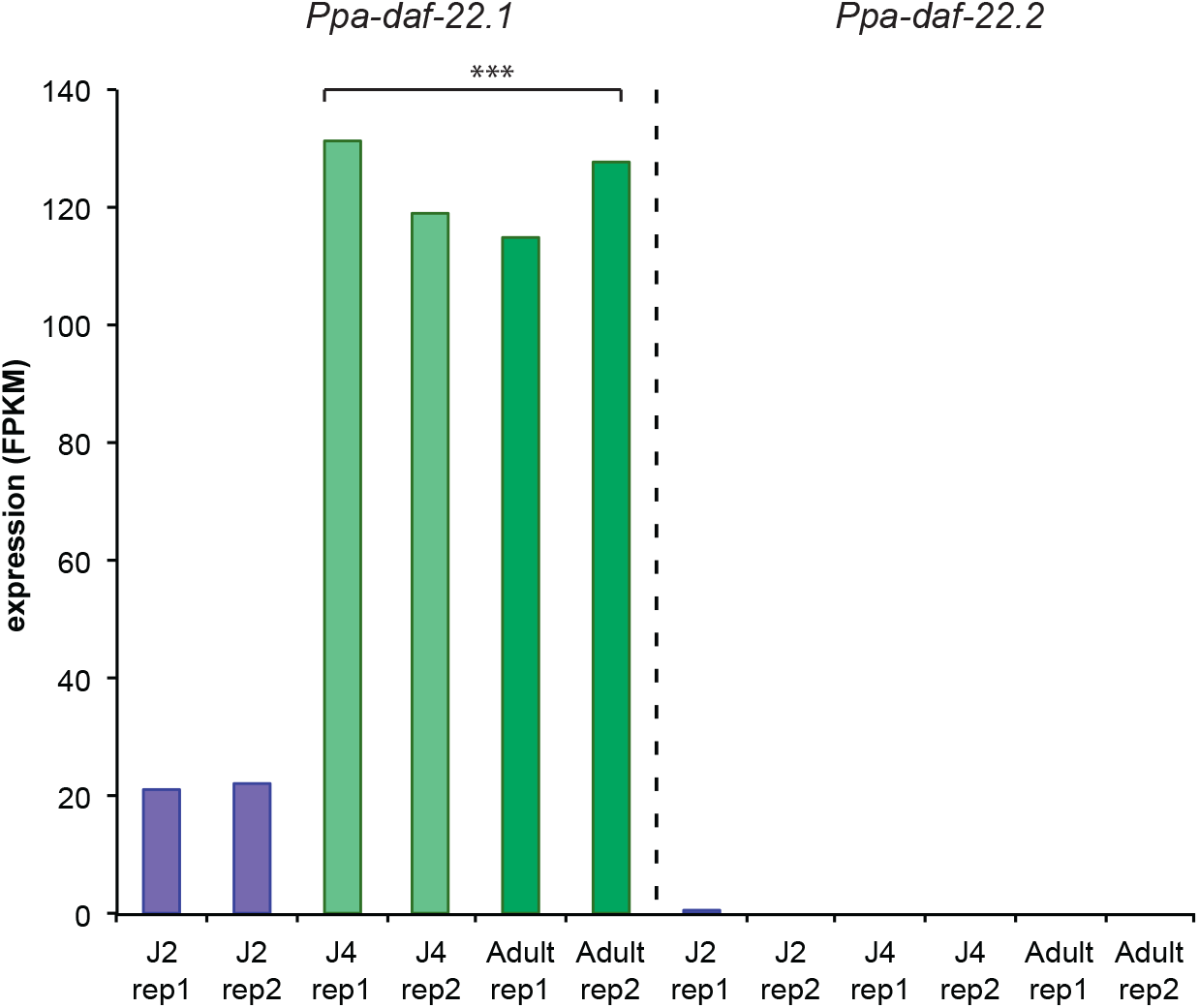
Enzyme that synthesize NDMMs is transcriptionally regulated during development. Comparison of *daf-22.1* and *daf-22.2* expression (FPKM) by RNA-seq through different stages of development, data from Baskaran et al., 2015. A two-sided students *t*-test was performed between J4-adults and J2s (Significance codes: 0 ‘***’ 0.001, ‘**’ 0.01, ‘*’ 0.05).

**Table S1, related to Figure 3.** Table of binomial regression *p* values for vital-dye method and excess crowding. Significance *p* values from binomial regression of vital-dye method for age and number added, and from binomial regression of number-reaching-adult and Eu counts for each number of individuals added relative to 1,000 individuals added.

**affect of population age on mouth form of developing juveniles**
| **binomial**<br>**regression** | ***p* value**<br>**J2s** | ***p* value**<br>**dauers** |
| --- | --- | --- |
| **age added** | 0.0259 | 0.002955 |
| **number added** | 4.28e-13 | 0.000404 |

**affect of number of peers on development and mouth form (proxy for potential starvation effects on mouth form)**
| **binomial regression** | ***p* value for development**<br>**(relative to 1,000)** | ***p* value for Eu**<br>**(relative to 1,000)** |
| --- | --- | --- |
| **3,000 J2s added** | 0.3408 | 1.0 |
| **4,000 J2s added** | 0.0424 | 1.0 |
| **5,000 J2s added** | 6.06E-14 | 0.99 |
| **10,000 J2s added** | 4.09E-14 | 0.99 |

**Table S2, related to Figure 5.** Table of linear regression *p* values with FDR corrections for strain and stage comparison of NDMM levels. FDR-corrected and uncorrected *p* values from linear regression of *P. pacificus* NDMMs (alternating grey background between NDMMs for clarity). Red values indicate FDR<0.05.

| NDMM comparison | pvalue | fdr corrected |
| --- | --- | --- |
| ascr1_stage | 0.4733 | 0.774490909 |
| ascr1_strain | 0.0429 | 0.110314286 |
| ascr1_stage:strain | 0.031 | 0.085846154 |
| ascr9_stage | 3.79E-05 | 0.0002274 |
| ascr9_strain | 0.651 | 0.778064516 |
| ascr9_stage:strain | 0.272 | 0.50148 |
| ascr12_stage | 0.0029 | 0.01404 |
| ascr12_strain | 0.0897 | 0.201825 |
| ascr12_stage:strain | 0.0302 | 0.085846154 |
| dasc1_stage | 9.62E-08 | 8.66E-07 |
| dasc1_strain | 0.11363 | 0.240628235 |
| dasc1_stage:strain | 0.00351 | 0.01404 |
| npar1_stage | 0.0033 | 0.01404 |
| npar1_strain | 0.9426 | 0.984 |
| npar1_stage:strain | 0.6355 | 0.778064516 |
| npar2_stage | 0.0516 | 0.12384 |
| npar_2strain | 0.984 | 0.984 |
| npar2_stage:strain | 0.9716 | 0.984 |
| pasc1_stage | 0.449 | 0.769714286 |
| pasc1_strain | 0.753 | 0.847125 |
| pasc1_stage:strain | 0.564 | 0.778064516 |
| pasc9_stage | 0.616 | 0.778064516 |
| pasc9_strain | 0.267 | 0.50148 |
| pasc9_stage:strain | 0.523 | 0.778064516 |
| pasc12_stage | 0.6122 | 0.778064516 |
| pasc12_strain | 0.2786 | 0.50148 |
| pasc12_stage:strain | 0.67 | 0.778064516 |
| tasc1_stage | 0.522 | 0.778064516 |
| tasc1_strain | 0.862 | 0.940363636 |
| tasc1_stage:strain | 0.57 | 0.778064516 |
| ubas1_stage | 3.13E-12 | 1.13E-10 |
| ubas1_strain | 0.00538 | 0.019368 |
| ubas1_stage:strain | 6.69E-08 | 8.03E-07 |
| ubas2_stage | 1.34E-11 | 2.41E-10 |
| ubas2_strain | 0.00711 | 0.023269091 |
| ubas2_stage:strain | 6.18E-07 | 4.45E-06 |

**Table S3, related to Figure 5.**
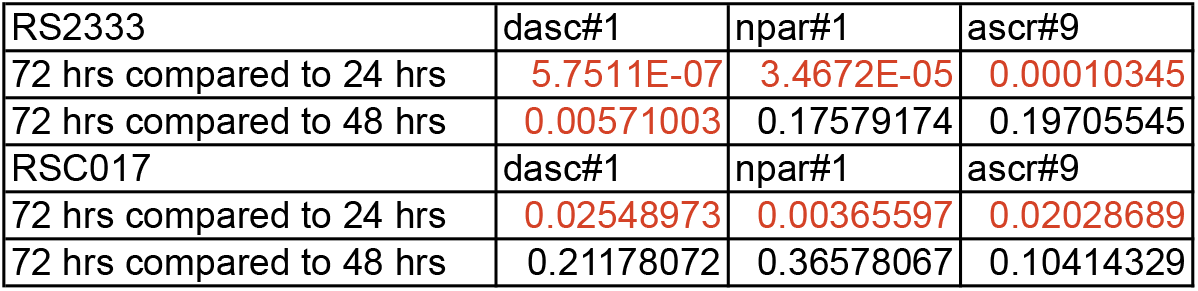
Pairwise comparison of dasc#1, npar#1, and ascr#9 throughout development. Significance assessed with a two-tailed student’s *t*-test. Red values indicate *p*<0.05.

